## Extended Data for "Assembly intermediates of orthoreovirus captured in the cell"

**Extended data figures 1-7 and tables 1-3.**

**Extended Data Fig. 1.** Maps coloured by radius for the 3 categories of particles, full and empty virion-like and stars (see Methods). The left hand panels show the whole particles, middle panels the particles are cropped at their centres in the z-direction, and the right hand panels show a thin slice at z=½. Note the density for the polymerase molecules in red in the empty virion-like particles.

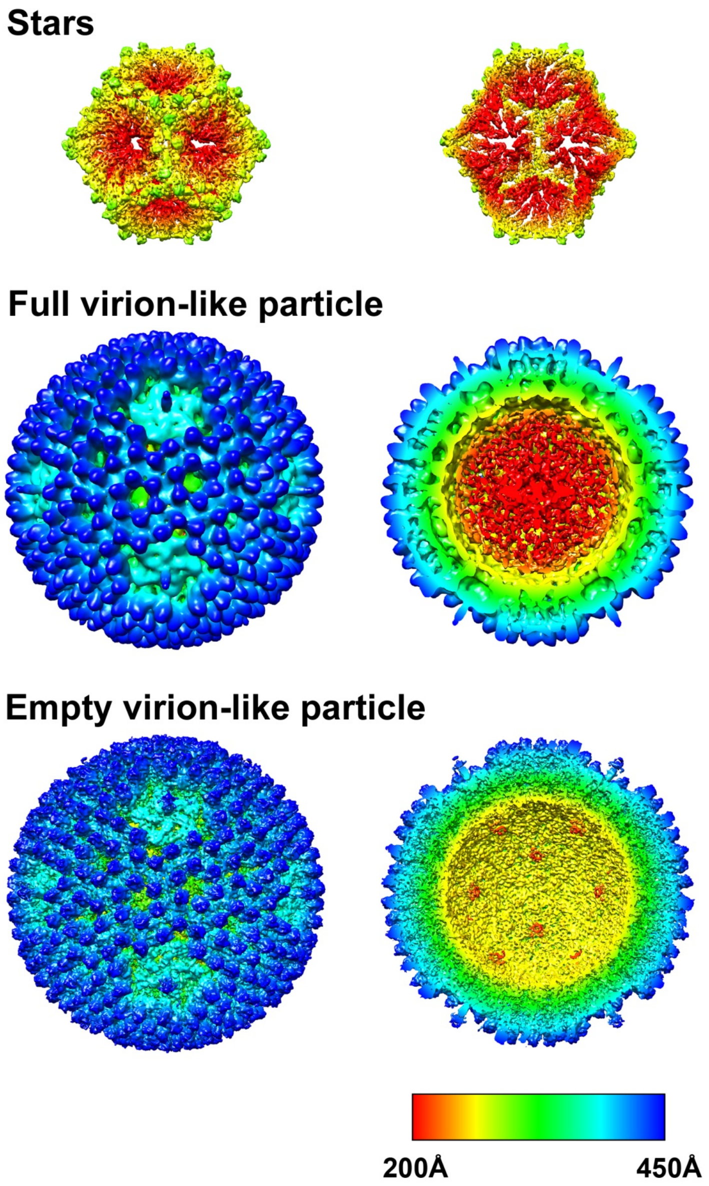

**Extended Data Fig. 2.** Sub-tomogram averaged maps from PEET for full (blue) and empty (red) virion-like particles. Note the density within the full particles (lower portion) and the common 5-fold spikes (centre top).

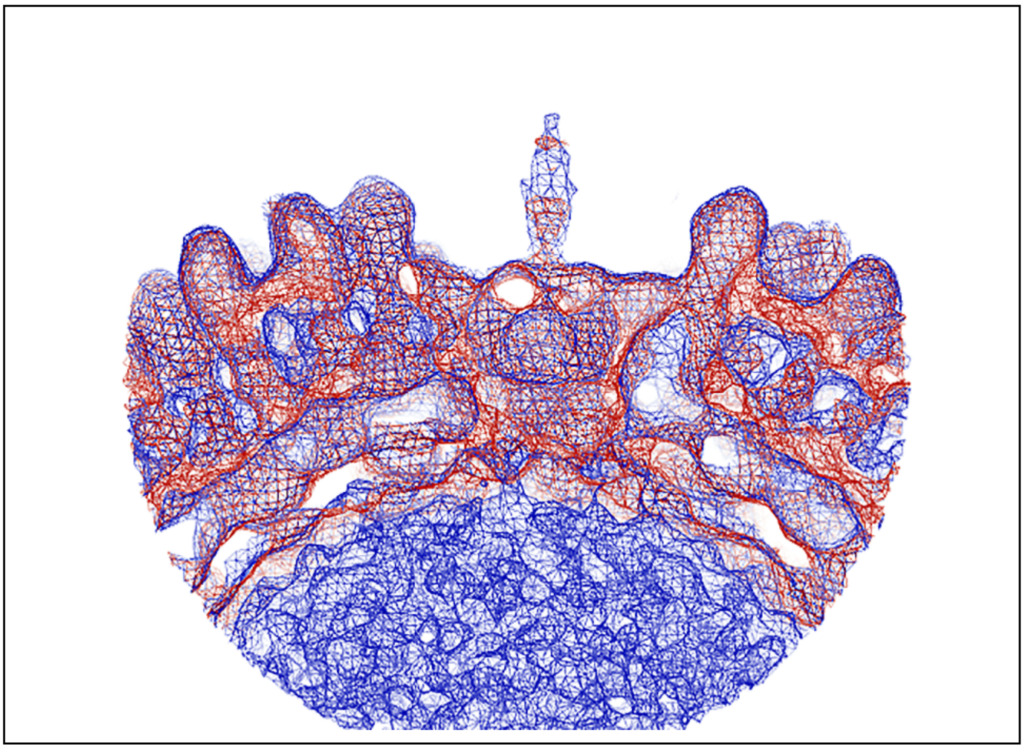

**Extended Data Fig. 3.** Definition of the axes used for local reconstructions in emClarity. Yellow shows pseudo 6-fold, red 5-fold.

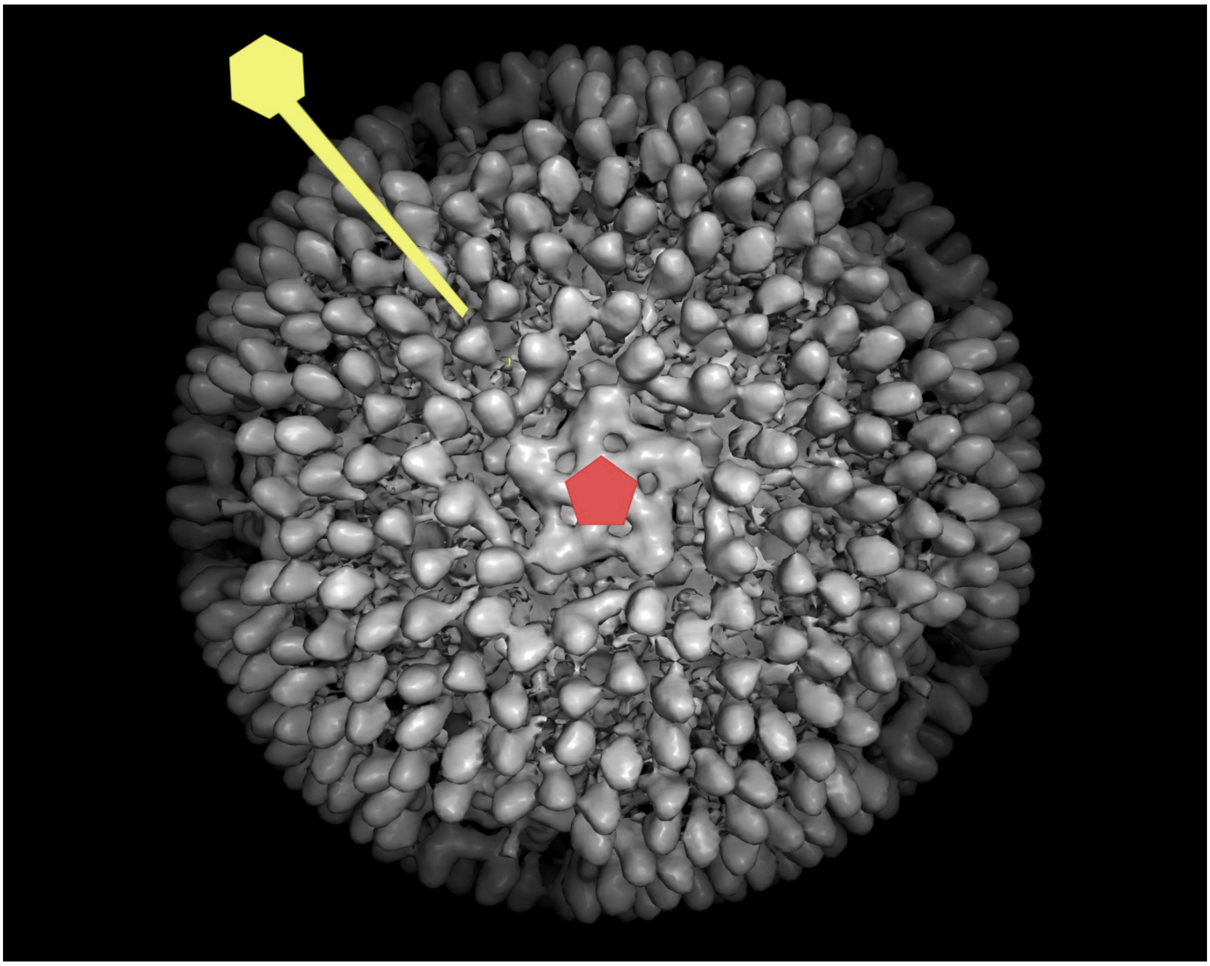

**Extended Data Fig. 4.** **a,** FSC curves for the five sub-icosahedral reconstructions. The FSC is between two half data sets. p6-fold corresponds to the pseudo 6-fold map. **b-d** Representative FSC curves for the fit of the models to the map (as calculated by the Phenix tool phenix.validation_cryoem[1,2]) for **b,** virion-like particle pseudo 6-fold, **c,** virion-like particle 5-fold and **d,** SLP. Masked data is coloured orange, unmasked blue.

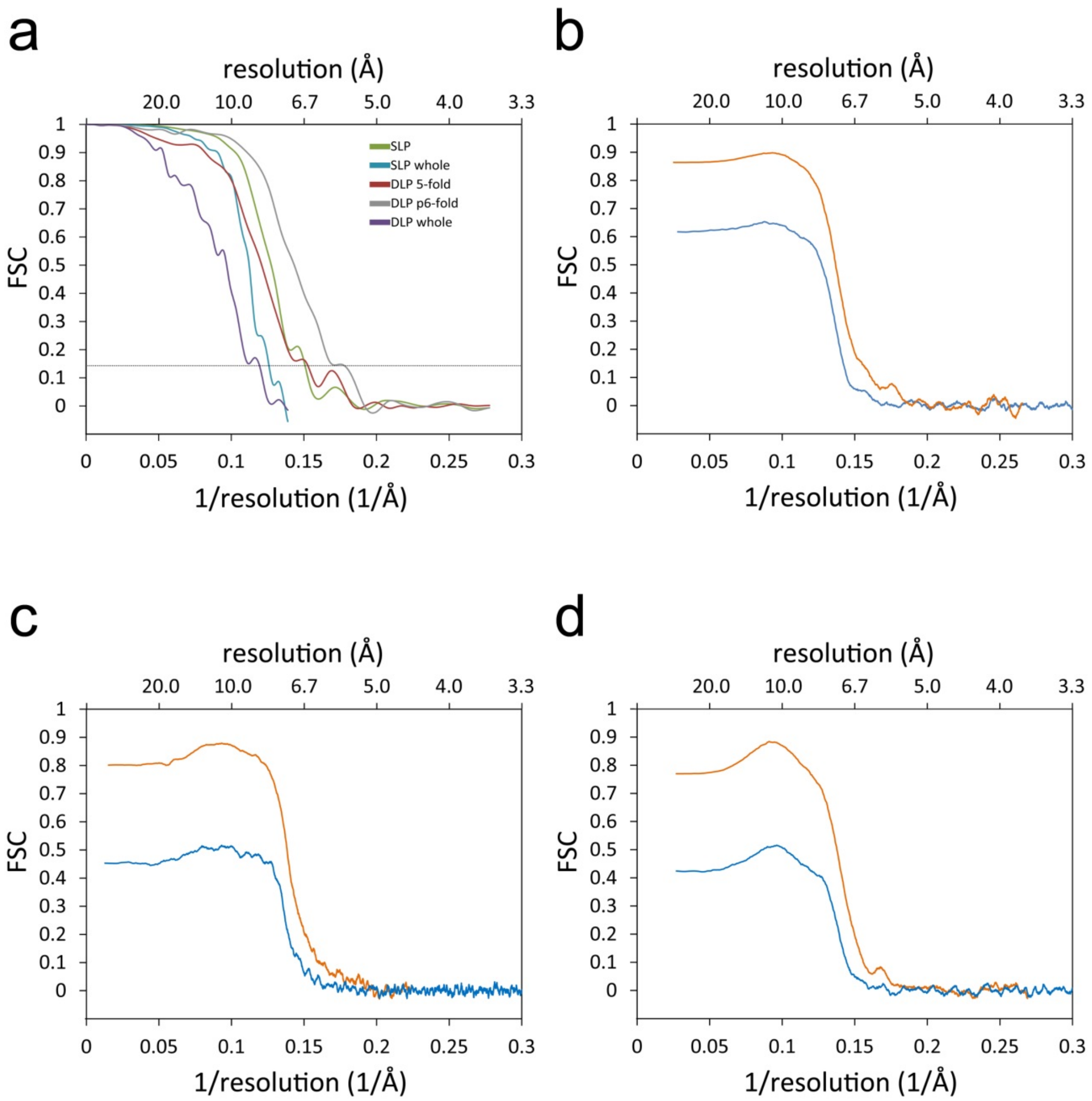

**Extended Data Fig. 5.** Rigid body fits for molecule A of **a,** reovirus λ1 and **b,** rotavirus VP2 into the SLP map. As can be seen there exists substantial density which cannot be accounted for by the rotavirus VP2. However, reovirus λ1 has insertions at residues 843-851 and 997-1014 which occupy this density.

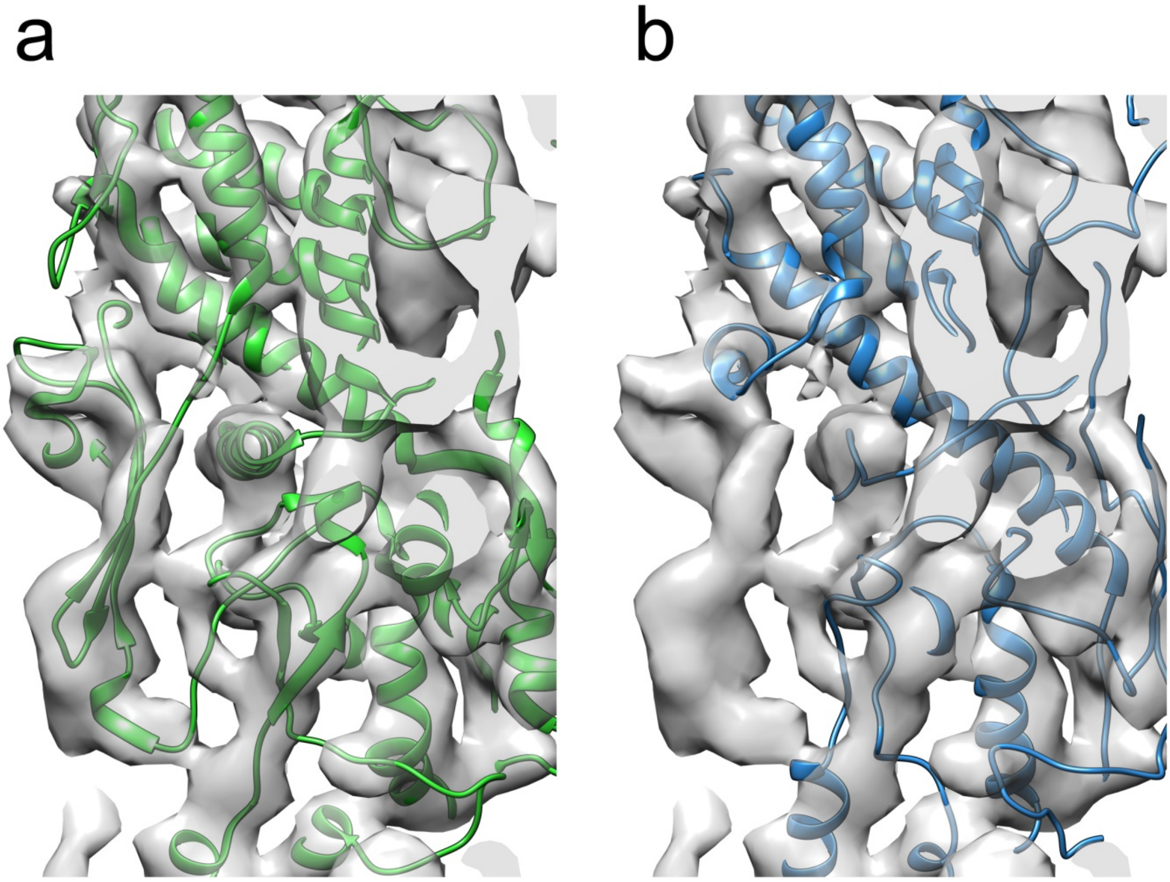

**Extended Data Fig. 6. Superposition of λ1 molecules from virion-like particles and SLP.** **a,** molecule A with virion-like particle coloured blue and SLP orange. **b,** molecule B coloured pale blue and red for virion-like particle and SLP, respectively.

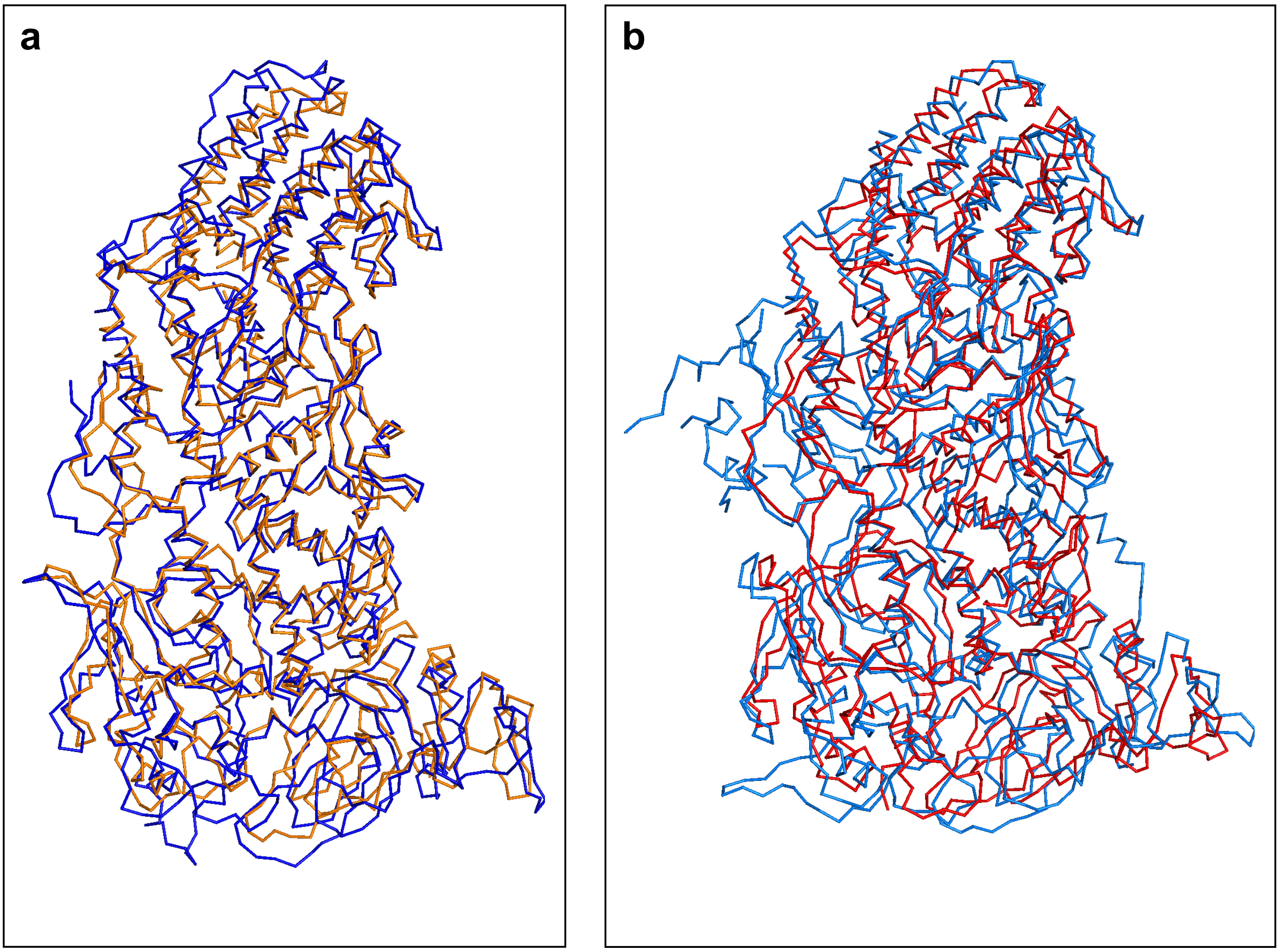

**Extended Data Fig. 7.** Roadmap figure showing the relative positions of the footprints for σ2 on the virion-like particle and the SLP. The λ1 molecules A and B are coloured faded green and red respectively. The footprint for the σ2 A-hinge is coloured blue, σ2 three-fold adjacent is purple and σ2 two-fold yellow. The roadmaps were produced using RIVEM[3].

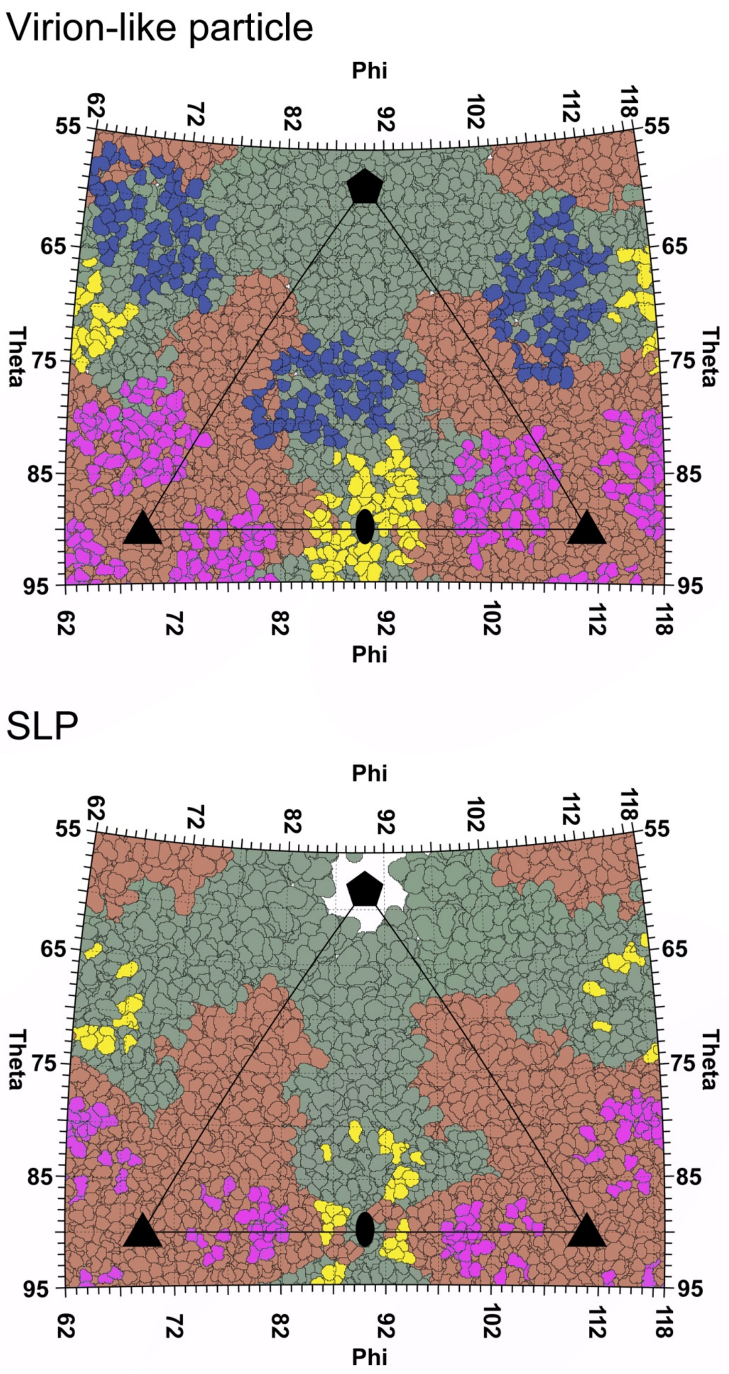

**Extended Data Table 1.** Summary of cryo-electron tomography data acquisition and image processing.

| | | | | | | |
|---|---|---|---|---|---|---|
| **Data Acquisition** | Microscope | Krios | | | | |
| | Voltage | 300 keV | | | | |
| | Detector | K2 Summit | | | | |
| | Energy filter | Yes, 20 eV slit | | | | |
| | Number of tomograms | 5 | | | | |
| | Defocus range | 4.3 to 5.9 | | | | |
| | Acquisition scheme | Dose-symmetric | | | | |
| | Total dose | 82 $e^{-}$/Å$^{2}$ | | | | |
| | Tilt range | -40° to +40° in 2° increments | | | | |
| **Data Processing** | Map | | | | | |
| | SLP-whole empty* | SLP 5-fold** | DLP-whole empty* | DLP 5-fold** | DLP pseudo 6-fold** | DLP- whole full* |
| Å/pixel | 3.6 | 1.8 | 3.6 | 1.8 | 1.8 | 3.6 |
| Number of sub-tomograms*** | 170 | 2,683 (242) | 18 | 625 (65) | 3,039 (65) | 10 |
| Final resolution | 7.8 Å | 6.6 Å | 8.3 Å | 6.5 Å | 5.6 Å | 17.0 Å |

* Icosahedral symmetry applied.

** Five- or six-fold symmetry applied.

*** In parentheses are the numbers of SLP or DLP particles included in sub-tomogram averaging of 5-fold and pseudo 6-fold maps.

**Extended Data Table 2.** Model refinement statistics.

| | Ramachandran plot (%) | | | Bonds (RMSD) | | Rotamer outliers (%) | Correlation coefficient | |
|---|---|---|---|---|---|---|---|---|
| Map and model | Favoured | Allowed | Outliers | Length (Å) | Angles(°) | | Main chain | Side chain |
| SLP | 86.40 | 13.50 | 0.10 | 0.998 | 0.006 | 0.40 | 0.79 | 0.77 |
| DLP 5-fold | 83.71 | 16.15 | 0.14 | 1.031 | 0.007 | 0.50 | 0.76 | 0.74 |
| DLP pseudo 6-fold | 90.00 | 9.93 | 0.07 | 0.921 | 0.006 | 0.31 | 0.78 | 0.77 |

**Extended Data Table 3.** Changes in the interface areas between λ1 molecules.

| Interface | Virion-like particle interface area (Å$^2$) | SLP interface area (Å$^2$) |
|---|---|---|
| A'B | 5,900 | 4,400 |
| AB | 5,500 | 4,000 |
| AB'' | 2,100 | 2,200 |
| BB' | 3,500 | 1,500 |
| AA' | 2,500 | 1,200 |
| AA'' | 1,200 | 1,000 |
